# An Aurora Kinase A/TPX2 complex phosphorylates CKAP2 to control mitotic spindle growth

**DOI:** 10.1101/2025.09.09.675182

**Authors:** Thomas J Kucharski, Tatiana Lyalina, Susanne Bechstedt

## Abstract

Faithful chromosome segregation requires the precise assembly of the mitotic spindle. Cytoskeleton-associated protein 2 (CKAP2) is a microtubule-associated protein with potent microtubule-polymerizing activity that localizes to spindle microtubules during mitosis. Loss or overexpression of CKAP2 causes chromosomal instability and aneuploidy, yet its regulation remains poorly understood. To identify CKAP2 interactors, we immunoprecipitated endogenous CKAP2 from mitotic RPE1 cells and analysed co-purifying proteins by mass spectrometry. This revealed a specific interaction between CKAP2 and the mitotic kinase Aurora A and its activator TPX2, but not with Aurora B, as previously suggested. We further show that CKAP2 co-localizes with Aurora A-TPX2 complexes throughout mitosis, and Aurora A directly phosphorylates CKAP2 in cells and in vitro. Phosphorylation of CKAP2 By Aurora kinase A decreases its microtubule affinity in cells and in vitro. These findings identify CKAP2 as a direct interactor of TPX2 and a substrate of Aurora kinase A and uncover a regulatory pathway controlling spindle growth and stability during mitosis.

## Introduction

The faithful segregation of chromosomes during mitosis is essential for maintaining genomic stability. At the onset of mitosis, the nuclear envelope breaks down and the mitotic spindle assembles to capture and segregate duplicated chromosomes^1^. The spindle is composed of microtubules, dynamic polymers of αβ-tubulin dimers that undergo continuous phases of growth and shrinkage. These dynamic properties generate the forces required to align chromosomes at the metaphase plate and to separate sister chromatids toward opposite poles once proper kinetochore attachments are established ^2,3^. Accurate chromosome segregation, therefore, depends on the precise spatial and temporal regulation of microtubule dynamics^4–7^, which is achieved through the coordinated actions of microtubule-associated proteins (MAPs)^1^. Among these MAPs, Cytoskeleton-Associated Protein 2 (CKAP2; also known as TMAP) has emerged as a potent microtubule polymerase that promotes microtubule growth and spindle assembly^8–14^. CKAP2 localizes to centrosomes and spindle microtubules during mitosis, and its overexpression leads to chromosomal instability, aberrant centrosome numbers, and poor prognosis in multiple cancers^8–14^. CKAP2 is phosphorylated on several residues during mitosis by Cyclin-dependent kinases (CDKs) and Aurora kinases^8,15–19^. Phosphorylation of conserved C-terminal sites has been proposed to regulate CKAP2 localization, which transitions from centrosomes and spindle poles during early mitosis to chromatin in late anaphase, followed by degradation via the Anaphase-Promoting Complex (APC) during telophase^8,15–20^. These observations suggest that reversible phosphorylation acts as a molecular switch controlling CKAP2 function and localization throughout mitosis.

Phosphorylation sites in the N terminus of CKAP2 have also been identified in large-scale phosphoproteomic datasets^21–23^, but their functional relevance remains unknown. To elucidate the molecular mechanisms regulating CKAP2 and identify possible kinases, we performed unbiased proteomic analyses to identify its interacting partners in mitotic cells. This approach revealed the mitotic kinase Aurora Kinase A (AurKA) and its cofactor, targeting protein for Xenopus kinesin-like protein 2 (TPX2) as prominent CKAP2-interacting proteins. We show that AurKA, in complex with TPX2, binds and phosphorylates CKAP2 during mitosis, thereby modulating its microtubule-binding and polymerase activities. These findings place CKAP2 within the AurKA-TPX2 signalling network and uncover a new regulatory pathway controlling spindle growth and mitotic fidelity.

## Results

### CKAP2 interacts with the AurKA-TPX2 complex during mitosis

To identify proteins that associate with CKAP2 during mitosis, we immunoprecipitated endogenous CKAP2 and its binding partners from RPE-1 cells synchronized in mitosis by sequential Palbociclib-STLC treatment (Fig. 1A). Mass spectrometry identified 672 proteins present in either CKAP2 or control immunoprecipitates. Applying stringent criteria (≥4 unique peptides and ≥4:1 enrichment over control) yielded a high-confidence set of 90 CKAP2-associated proteins (Supplemental Table 1). Gene ontology analysis of this dataset revealed strong enrichment for proteins involved in mitosis, chromatin organization, and RNA processing (Fig. 1B). Interaction network mapping (STRING) further linked CKAP2 to functional clusters associated with spindle organization. Among MAPs, KIF2A, KIF20A, KIF23, TXLNA, and TPX2 were identified, together with AurKA, a well-known mitotic kinase and binding partner of TPX2^24–31^. The dataset also contained chromatin-associated proteins, including TOP2A, SMC2/4, NCAPG/D2/H, MACROH2A1, and KI67, that might mediate the transition of CKAP2 to chromatin in late mitosis (Fig. 1C, 1D).

**Figure 1:**
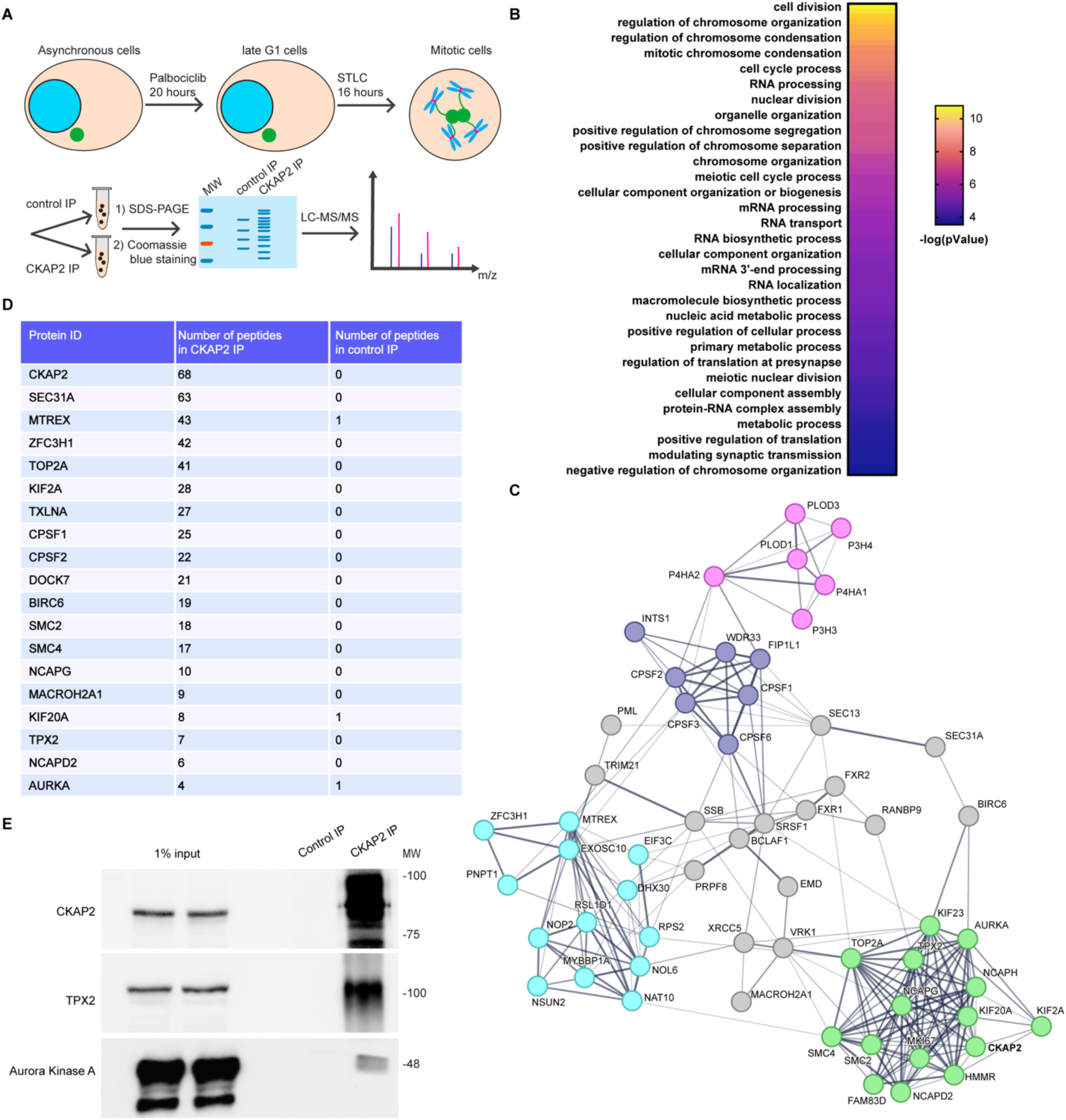
Proteomic screening identifies AurKA/TPX2 as CKAP2-interacting proteins. A) Cartoon diagram showing the experimental scheme for determining CKAP2 binding partners. RPE1 cells were first synchronized in mitosis by Palbociclib-STLC block. The cells were then harvested and lysed. Endogenous CKAP2 was then immunoprecipitated from the mitotic lysate together with interacting proteins. The purified proteins were then separated via SDS-PAGE. The gel lanes were then cut into 12 slices and analyzed by mass spectrometry to identify CKAP2 interaction partners. A single experiment was performed. B) Proteins from the phospho-proteomic screen from (A) that that met criteria of at least 4 peptides detected and 4:1 signal to noise ratio were analyzed for gene ontology. The data were then transferred to Prism software and plotted. C) STRING analysis of the proteins from (A) meeting detection criteria showing protein interaction networks. D) Selected interacting proteins detected in the control immunoprecipitation and CKAP2 immunoprecipitations from (A) showing the number of total detected peptides. E) RPE1 cells were prepared as in panel (A), except that purified proteins were analyzed by Western blot. CKAP2 was immunoprecipitated from Palbociclib-STLC mitotic lysates and the purified proteins were then separated by SDS-PAGE, transferred to nitrocellulose membrane and blotted as indicated. The panels were adjusted for brightness and contrast. An experiment representative of two independent biological repeats is shown.

Given the central role of AurKA and TPX2 in spindle regulation, we focused on their interaction with CKAP2. AurKA phosphorylates multiple substrates at spindle poles and microtubules to promote spindle assembly and stability^24–27^, whereas TPX2 binds directly to microtubules and recruits AurKA to the spindle, activating the kinase locally^28,32,33^. Because TPX2 is essential for the localization and activity of AurKA^28^, and, like CKAP2, both are frequently overexpressed in cancer^30,34,35^, we hypothesized that CKAP2 may function within this regulatory network ^32,33,36^.

To validate the CKAP2-AurKA interaction biochemically, we immunoprecipitated CKAP2 from mitotic RPE-1 cells and probed for AurKA and TPX2 by immunoblotting. Both proteins robustly co-precipitated with CKAP2 (Fig. 1E). Reciprocal immunoprecipitation of TPX2 confirmed its association with both AurKA and CKAP2 (Fig. S1A). AurKA shares high sequence and substrate similarity with Aurora B kinase (AurKB), yet the two are spatially segregated during mitosis: TPX2 localizes AurKA to spindle poles and microtubules, whereas INCENP recruits AurKB to centromeres and the midbody^24–27,37–39^. AurKB has previously been proposed to phosphorylate CKAP2^19^. Therefore, we compared CKAP2 localization relative to both kinases. Using RPE-1 cells with endogenous GFP-labelled CKAP2 (knock-in)^8^, we co-stained for total AurKA and its active, phosphorylated form (pT288)^29,31^. CKAP2 strongly co-localized with AurKA and pT288-AurKA at centrosomes in prophase and along pole-proximal spindle microtubules during prometaphase and metaphase (Fig. 2A). Similarly, CKAP2 extensively co-localized with TPX2 in these regions (Fig. 2B), consistent with our proteomic and biochemical data (Fig. 1; Supplemental Table 1). In contrast, AurKB localized to centromeres during prophase-metaphase and to the midbody during anaphase, with no detectable overlap with CKAP2 (Fig. S1B-C). Treatment of cells with the AurKB inhibitor ZM447439, at doses sufficient to cause severe segregation defects, did not alter CKAP2 localization on the spindle or chromatin (Fig. S1D). Together, these data demonstrate that CKAP2 specifically associates and co-localizes with the AurKA-TPX2 complex, but not with AurKB, during mitosis.

**Figure 2:**
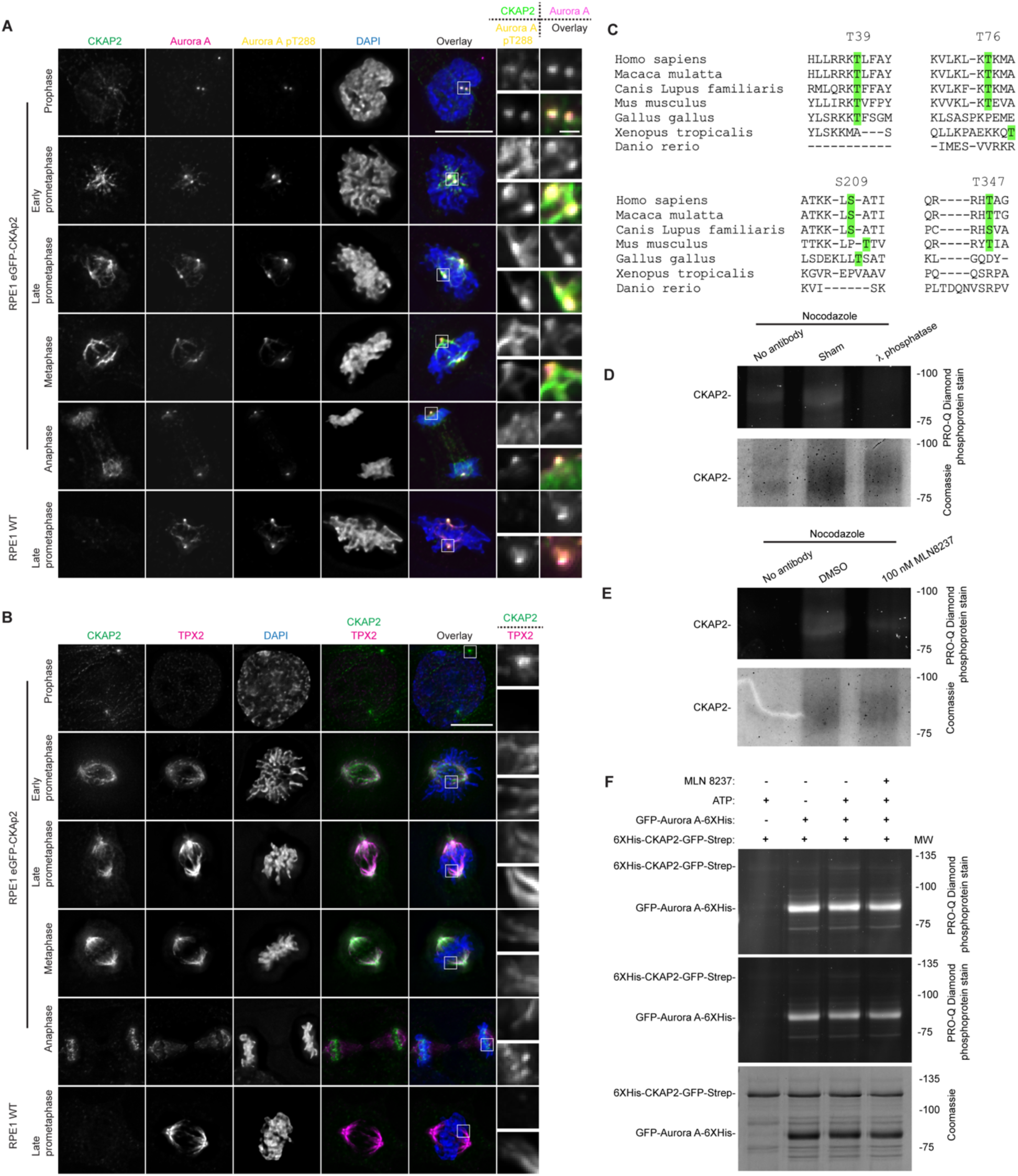
CKAP2 co-localizes with and is phosphorylated by AurKA/TPX2. A) Immunofluorescence images from RPE1 eGFP-CKAP2 cells in various stages of mitosis stained for AurKA, AurKA pT288 and DAPI. The CKAP2-eGFP, AurKA and AurKA pT288 channels were adjusted evenly for brightness and contrast. The DAPI channel of each condition was adjusted independently. Representative images from 2 independent biological experiments are shown. Scale bars, 10 μm and 1 μm (inset). B) Immunofluorescence images from RPE1 eGFP-CKAP2 cells in various stages of mitosis stained for TPX2 and DAPI. The CKAP2-eGFP and TPX2 channels were adjusted evenly for brightness and contrast. The DAPI channel of each condition was adjusted independently. Representative images from 2 independent biological experiments are shown. Scale bars, 10 μm and 1 μm (inset). C) Amino acid sequence alignment of CKAP2 orthologs showing predicted AurKA phosphorylation consensus motifs. Residues highlighted in green indicate potential AurK phospho-acceptor residues. D) RPE1 cells were synchronized by Palbociclib-STLC block. The cells were then harvested and lysed by boiling in SDS-containing buffer. The SDS was then inactivated by 10-fold dilution with 1.5% Triton X-100 buffer. CKAP2 was then purified by immunoprecipitation. The protein-G agarose beads were either sham treated or treated with Lambda protein phosphatase for 30 minutes at 30°C. The beads were then boiled in sample buffer to elute purified proteins. Purified proteins were then separated by SDS-PAGE, which was then stained with ProQ Diamond phosphoprotein stain and imaged. Finally, the gel was stained with Coomassie blue and imaged again. An experiment representative of two independent biological repeats is shown. E) Cells were prepared as in (D), except that one set of cells was treated concurrently with STLC and DMSO control and one set with MLN 8237 AurKA inhibitor. CKAP2 was then purified and analyzed as in (D). An experiment representative of two independent biological repeats is shown. F) Purified GFP-AurKA-6XHis and His-mmCKAP2-GFP-Strep were mixed with reaction components as indicated and incubated at 37°C for 1 hr. The reaction was then stopped by addition of 4X sample buffer, boiled and separated by SDS-PAGE gel. The gel was then stained with ProQ Diamond phosphoprotein stain, imaged and subsequently stained with Coomassie blue and re-imaged. An experiment representative of two independent biological repeats is shown. High-brightness/contrast and low-brightness/contrast images are shown for presentation.

### AurKA-TPX2 phosphorylates CKAP2 during mitosis

To determine whether CKAP2 is a substrate of AurKA, we first analyzed its primary sequence for AurKA consensus motifs (R/K-R/K-x-S/T). Putative phosphorylation sites were identified at residues Ser39, Ser76, Thr209, and Ser347 (corresponding to mouse CKAP2 residues Ser52, Ser86, Thr206, and Ser337), in addition to several previously reported C-terminal sites^16–18^. These N-terminal sites are conserved across mammals but not in more distant species (Fig. 2C). Examination of published phosphoproteomic datasets confirmed phosphorylation at Thr209, supporting the physiological relevance of this site ^21–23^.

To test whether CKAP2 is phosphorylated during mitosis, we immunoprecipitated endogenous CKAP2 from RPE-1 cells synchronized by Palbociclib-STLC block. Samples were boiled in buffer containing SDS to ensure recovery of both soluble and microtubule-bound CKAP2. Treatment of the immunoprecipitate with λ-protein phosphatase caused a complete loss of the ProQ Diamond phosphorylation signal, confirming that CKAP2 is phosphorylated in mitotic cells (Fig. 2D).

We next asked whether AurKA is responsible for this phosphorylation. Mitotic RPE-1 cells were treated with either DMSO or the AurKA inhibitor MLN8237 prior to lysis and CKAP2 purification. Phosphoprotein staining revealed a marked decrease in CKAP2 phosphorylation upon MLN8237 treatment, indicating that AurKA activity is a major determinant of CKAP2 phosphorylation in mitosis (Fig. 2E). To test this directly, we expressed and purified CKAP2-GFP from E. coli and incubated the protein *in vitro* with recombinant GFP-AurKA (Fig. S1E) in the presence or absence of ATP. CKAP2 phosphorylation was readily detected only when active AurKA and ATP were included, demonstrating that AurKA directly phosphorylates CKAP2 in a kinase-dependent manner (Fig. 2F). Together, these findings establish that CKAP2 binds, co-localizes with, and is phosphorylated by the AurKA-TPX2 complex during mitosis.

### CKAP2 phosphorylation modulates its microtubule-binding affinity

Phosphorylation commonly regulates the interaction of MAPs with microtubules by introducing negative charges that weaken electrostatic interactions with the negatively charged tubulin surface. For example, phosphorylation of the kinetochore protein Hec1 at multiple sites fine-tunes its microtubule-binding affinity^40–43^. We therefore asked whether AurKA-dependent phosphorylation of CKAP2 similarly affects its association with spindle microtubules.

To test this in cells, we treated RPE-1 cells with 200 nM MLN8237 to inhibit AurKA and examined CKAP2 localization relative to the mitotic spindle. AurKA inhibition produced a clear increase in CKAP2 overlap with spindle microtubules. While total CKAP2 fluorescence intensity remained essentially unchanged, β-tubulin signal intensity on the spindle decreased by approximately 50%, resulting in an overall twofold increase in normalized CKAP2 association with microtubules (Fig. 3A-E). These findings suggest that AurKA-mediated phosphorylation negatively regulates CKAP2’s microtubule binding in cells.

**Figure 3:**
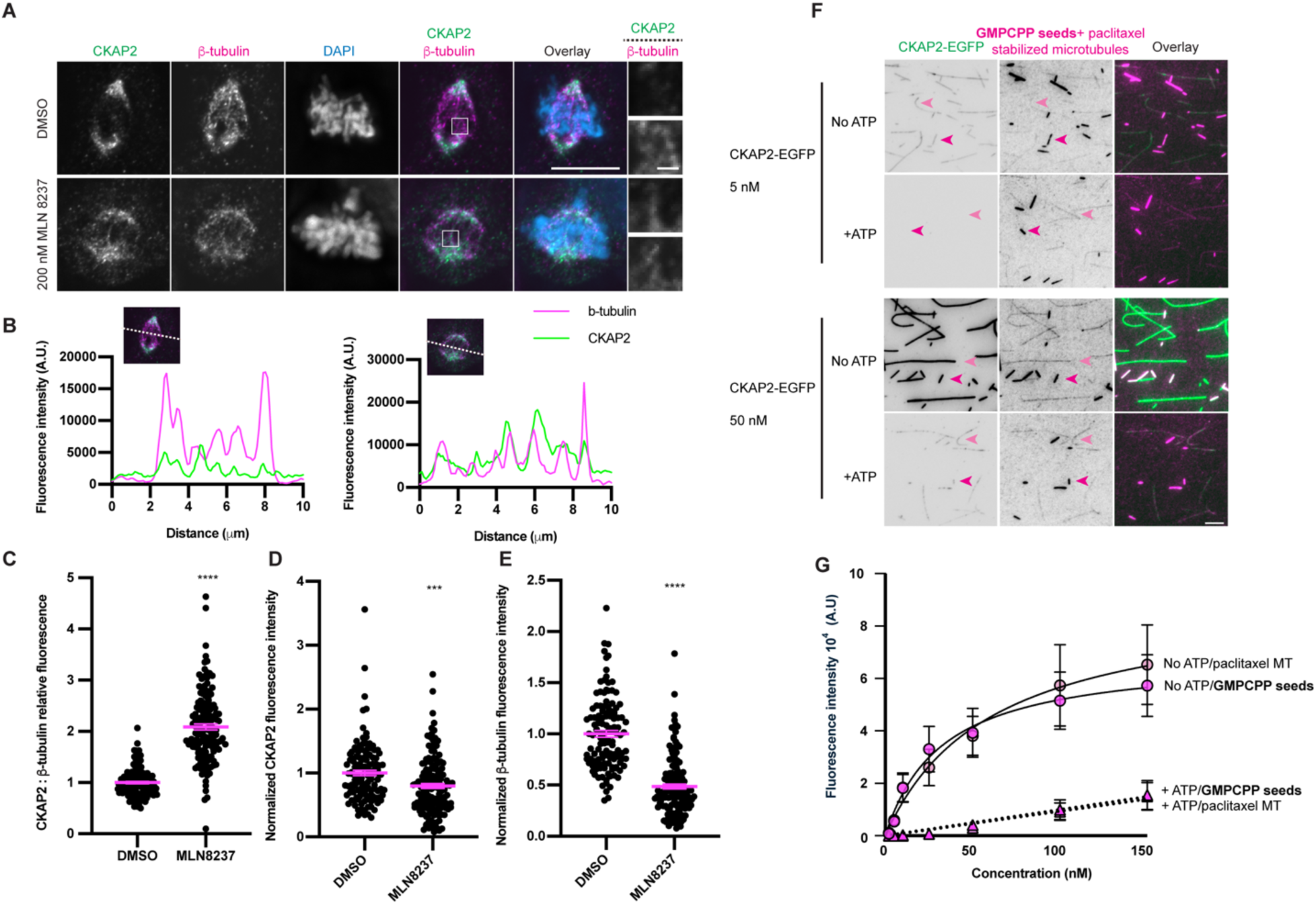
Phosphorylation by AuroraA reduces the mitotic spindle localization of CKAP2. A) Immunofluorescence images from asynchronous RPE1 cells in late prometaphase were treated with either DMSO or MLN8237 for 3 hours and then fixed and stained for CKAP2, β -tubulin and DAPI. The CKAP2 and β -tubulin channels were adjusted evenly for brightness and contrast. The DAPI channel of each condition was adjusted independently. Representative images from 2 independent biological experiments are shown. Scale bars, 10 μm and 1 μm (inset). B) Line scans across the indicated paths from the images in panel (A). C) Quantification of the CKAP2-β-tubulin ratio from panel (A). The Z-section with the brightest signal was chosen, and an outline was traced around the mitotic spindle. The levels of CKAP2 and β -tubulin were then measured. The background signal was taken as the minimum measured value in the mitotic spindle area and subtracted from the measured signal. At least 50 cells per condition for each of 3 independent biological repeats were measured. Error bars indicate the mean +/- SEM. Statistical significance was calculated between the indicated conditions using Student’s T- test. D) Quantification of the CKAP2 levels from panel (A) as for panel (C). E) Quantification of the β -tubulin levels from panel (A) as for panel (C). F) TIRF microscopy images of CKAP2 together with GMPCPP- and paclitaxel-stabilized microtubules. CKAP2 was incubated *in vitro* with AurKA in the presence or absence of ATP. Channels in between silanized-glass coverslips were prepared by flowing in anti-TAMRA antibodies. After washing, TAMRA labelled GMPCPP- and paclitaxel-stabilized microtubules were flowed into the chamber. Then, CKAP2, prepared as indicated, was also flowed into the chamber, and images were taken. Representative images from 2 independent biological experiments are shown. Scale bar, 5 μm. G) Quantification of the results presented in (F). The fluorescence intensity of CKAP2 on each of 20 GMPCPP -or 20 paclitaxel-stabilized microtubules was quantified.

To determine whether this effect is direct, we used total internal reflection fluorescence (TIRF) microscopy to visualize interactions between purified CKAP2 and stabilized microtubules *in vitro*. Recombinant CKAP2 was incubated with AurKA in the presence or absence of ATP to generate phosphorylated or unphosphorylated CKAP2, respectively, as in our biochemical assay (Fig. 2F). Each preparation was then incubated with GMPCPP-and paclitaxel-stabilized microtubules. Phosphorylated CKAP2 displayed a profound loss of microtubule binding: at 5 nM concentration, unphosphorylated CKAP2 bound readily along microtubule lattices, whereas phosphorylated CKAP2 showed no detectable binding. At 50 nM, phosphorylated CKAP2 bound weakly but still at 6-fold less compared the unphosphorylated protein (Fig. 3F-G). Together, these data demonstrate that AurKA-dependent phosphorylation of CKAP2 directly reduces its affinity for microtubules, providing a mechanism by which spindle-associated kinase activity modulates CKAP2 function during mitosis.

### TPX2 facilitates CKAP2 phosphorylation by AurKA

Given that CKAP2 associates with the AurKA-TPX2 complex (Fig. 1C-E; Fig. S1A), we next asked whether CKAP2 contributes to TPX2 localization on the mitotic spindle. To test this, we compared TPX2 fluorescence intensity on spindles in CKAP2 knockout (KO) RPE-1 cells^8^ and wild-type controls. TPX2 spindle levels were significantly decreased in two of three CKAP2 KO clones (Fig. 4A, 4B). All three CKAP2 KO clones also exhibited reduced tubulin intensity (Fig. 4A, 4 C), suggesting that CKAP2 may either directly promote TPX2 recruitment to spindle microtubules or influence TPX2 levels indirectly by modulating overall microtubule levels.

**Figure 4:**
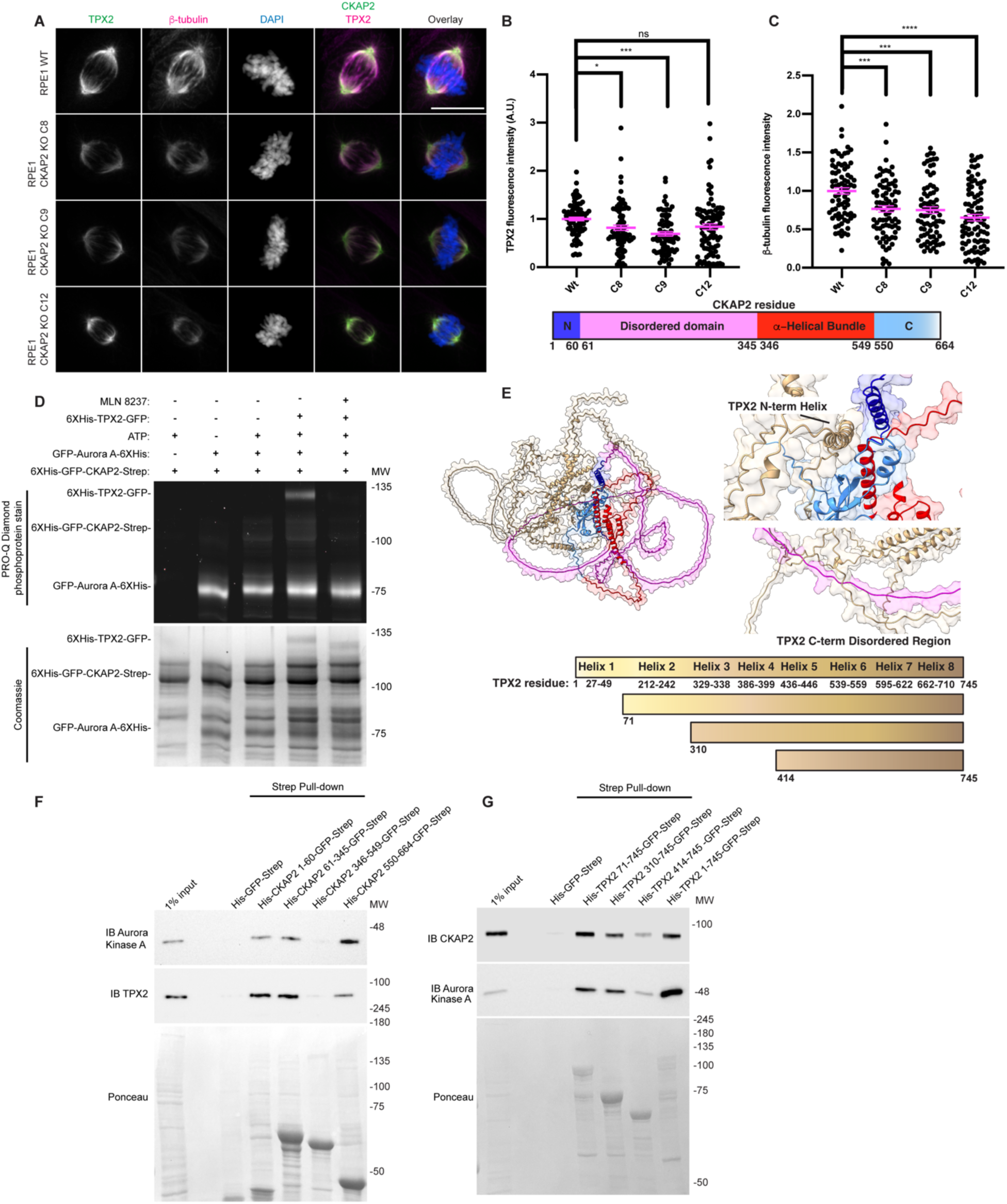
TPX2 and CKAP2 reciprocally regulate their microtubule binding activities. A) Immunofluorescence images of asynchronous RPE1 wild-type and CKAP2 KO clones 8, 9 and 12. The cells were plated on glass coverslips, allowed to proliferate and then fixed and stained for TPX2, β−tubulin and DAPI. The CKAP2 and β-tubulin channels were adjusted evenly for brightness and contrast. The DAPI channel of each condition was adjusted independently. Representative images from 2 independent biological experiments are shown. Scale bars, 10 μm. B) Quantification of the TPX2 levels from panel (A). The Z-section with the brightest signal was chosen, and an outline traced around the mitotic spindle. The levels of TPX2 were then measured. Background signal was taken as the minimum measured value in the mitotic spindle area and subtracted from the measured signal. At least 25 cells per condition for each of 3 independent biological repeats were measured. Error bars indicate the mean +/- SEM. Statistical significance was calculated between the indicated conditions using Dunnett’s multiple comparison test. C) As for panel (B) but the β−tubulin levels were quantified D) Purified GFP-AurKA-6XHis, 6XHis-mmTPX2-GFP-Strep and 6XHis-mmCKAP2-GFP-Strep were mixed with reaction components as indicated and incubated at 37°C for 1 hr. The reaction was then stopped by addition of 4X sample buffer, boiled and separated by SDS-PAGE gel. The gel was then stained with ProQ Diamond phosphoprotein stain, imaged and subsequently stained with Coomassie blue and re-imaged. The images were adjusted for brightness and contrast for presentation. An experiment representative of two independent biological repeats is shown. E) Alphafold2 model of mmTPX2 (gold) bound to mmCKAP2 (coloured by domain), highlighting the interaction between the TPX2 alpha helix 1 and the CKAP2 n-terminal helix and C-terminus domain. The domain structures of CKAP2 and TPX2, based on Alphafold models, are also shown. F) 6XHis-mmCKAP2-GFP-Strep fragments were purified by one-step Strep affinity gel purification. Concurrently, HeLa cells were synchronized in mitosis by thymidine-nocodazole block. The cells were then harvested and lysed. The lysates were then added to the Strep affinity gel bound to mmCKAP2 fragments and incubated. Following washes, the purified proteins were then separated by SDS-PAGE and transferred to a nitrocellulose membrane and finally stained with Ponceau and blotted as indicated. The images were adjusted for brightness and contrast for presentation. An experiment representative of two independent biological repeats is shown. G) 6xHis-mmTPX2-GFP-Strep deletion mutants were purified by one-step Strep affinity gel purification. Concurrently, HeLa cells were synchronized in mitosis by thymidine-nocodazole block. The cells were then harvested and lysed. The lysates were then added to the Strep affinity gel bound to mmTPX2 deletion mutants and incubated. The purified proteins were then separated by SDS-PAGE and transferred to a nitrocellulose membrane and finally stained with Ponceau and blotted as indicated. The images were adjusted for brightness and contrast for presentation. An experiment representative of two independent biological repeats is shown.

Our proteomic and immunoprecipitation analyses indicated that CKAP2 interacts directly with TPX2 and, to a lesser extent, with AurKA, consistent with higher TPX2 peptide abundance in mass spectrometry and a stronger TPX2 signal in immunoblots. We therefore asked whether TPX2 enhances CKAP2 phosphorylation by serving as a bridging factor for AurKA. In *in vitro* kinase assays performed with or without TPX2 (Fig. S1E), the presence of TPX2 markedly increased CKAP2 phosphorylation by AurKA (Fig. 4D). As expected, TPX2 itself was efficiently phosphorylated by AurKA in the same reaction. These results suggest that TPX2 not only recruits AurKA to CKAP2 but also enhances its phosphorylation efficiency.

To determine how CKAP2 and TPX2 may interact, we modelled the CKAP2-TPX2 complex using AlphaFold2^44^. The predicted complex revealed extensive electrostatic contacts between the basic residues of the CKAP2 N-terminal α-helix, several residues within its disordered region, and mixed-charge residues within the C-terminal domain, all of which engage TPX2 (Fig. 4E, Supplementary Movie 1). To definitively map the interaction domains, we performed pull-down assays using fragments of mouse CKAP2 encompassing the N-terminal α-helical region (1-60), the disordered central domain (61-345), the α-helical bundle (346-549), and the C-terminal domain (550-664^45^. The disordered domain bound TPX2 and AurKA most strongly, while the N-terminal and C-terminal regions also showed detectable, though weaker, interactions. The α-helical bundle region failed to bind either protein (Fig. 4E and F).

The AlphaFold model predicts that CKAP2 primarily interacts with the N-terminal α-helix 1 region of TPX2. To corroborate these findings, we next examined which regions of TPX2 mediate its interaction with CKAP2 and AurKA. Using TPX2 deletion mutants as bait, we found that removal of the N-terminal α-helix 1 dramatically reduced binding to both CKAP2 and AurKA, consistent with prior reports that this region is critical for AurKA association^28^. Progressive truncations of TPX2 further weakened CKAP2 binding, suggesting multiple contact sites between the proteins (Fig. 4E and G). Together, these findings demonstrate that CKAP2 and TPX2 depend upon discrete structural elements for binding and that they might function together in a complex to regulate microtubules.

### A CKAP2-AurKA-TPX2 complex regulates microtubule growth and stability

Given that CKAP2 and TPX2 physically interact and both affect microtubule behaviour^8,9,32,36,46–49^, we next tested if feedback from microtubules could affect AurKA phosphorylation activity towards CKAP2. Thus, we added increasing quantities of paclitaxel-stabilized microtubule s to an *in vitro* kinase assay. Interestingly, the addition of paclitaxel-stabilized microtubules increased the phosphorylation of CKAP2 by AurKA in a dose-dependent manner (Fig. 5A). The presence of microtubules strongly promoted phosphorylation of CKAP2 but did not affect phosphorylation of TPX2, likely because AurKA and TPX2 rapidly dimerize, while the microtubules promote complex formation between AurKA/TPX2 and CKAP2.

**Figure 5:**
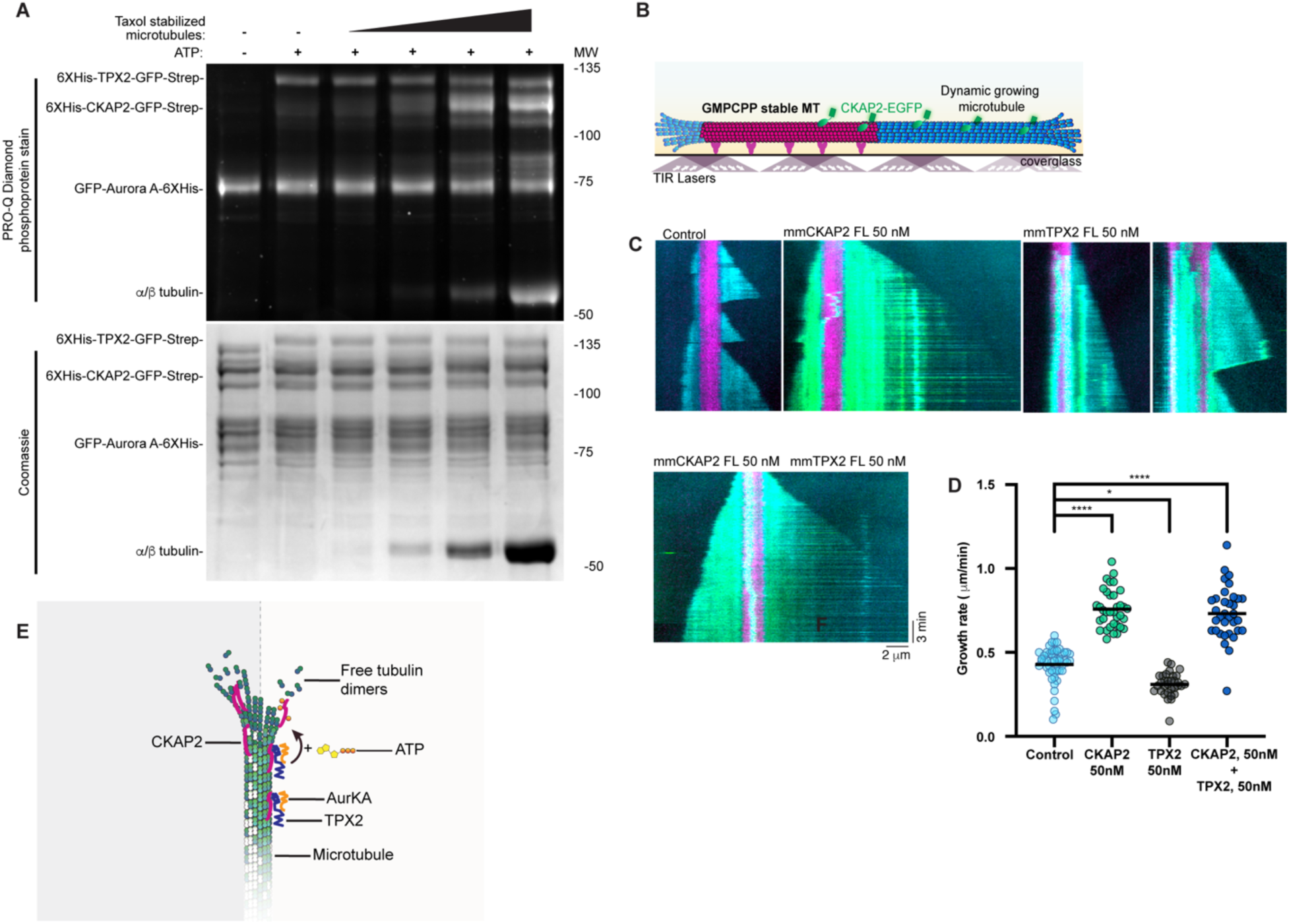
A CKAP2-AurKA-TPX2 complex regulates microtubule functions. A) Purified GFP-AurKA-6XHis, 6XHis-mmTPX2-GFP-Strep and 6XHis-mmCKAP2-GFP-Strep were mixed with increasing amounts of paclitaxel-stabilized microtubules and incubated at 37 °C for 1 hr. The reaction was then stopped by the addition of 4X sample buffer, boiled and separated by SDS-PAGE gel. The gel was then stained with ProQ Diamond phosphoprotein stain, imaged and subsequently stained with Coomassie blue and re-imaged. The images were adjusted for brightness and contrast for presentation. An experiment representative of two independent biological repeats is shown. B) Cartoon diagram showing the setup of the dynamic microtubule growth assay utilized in (G). C) Kymographs of TIRF microscopy images of His-mmCKAP2-GFP-Strep and/or His-mmTPX2-GFP-Strep together with dynamic microtubules. GMPCPP seeds were flowed into a slide chamber and washed. Free tubulin and GTP were then flowed into the chamber along with His-mmCKAP2-eGFP-Strep, His-mmTPX2-eGFP-Strep or both proteins together. Images were then taken at the indicated time points. Representative images of one of two independent biological repeats are shown. D) Quantification of the data from (D). The growth rate from at 51 (control) 35 (CKAP2) 32 (TPX2) 34 (CKAP2+TPX2) microtubules per condition from two independent experiments were measured and plotted in Prism. E) Cartoon diagram showing how CKAP2 is affected by phosphorylation via an AurKA/TPX2 complex. By itself, CKAP2 robustly mediates microtubule growth and stabilization. During mitosis, CKAP2 is bound by TPX2, which results in phosphorylation via the TPX2 binding partner AurKA. Phosphorylation of CKAP2 results in lower microtubule binding affinity and polymerization activity.

Finally, we tested whether CKAP2 and TPX2 act cooperatively to modulate microtubule dynamics. Using an *in vitro* microtubule polymerization assay, we compared the effects of purified CKAP2 and TPX2 individually and in combination. In control reactions lacking either protein, microtubules exhibited dynamic growth and frequent depolymerization. As reported previously, 50 nM CKAP2 strongly enhanced microtubule growth and suppressed catastrophe events^9^, while 50 nM TPX2 alone slightly stabilized microtubules and reduced polymerization rates^36^. When both proteins were combined, CKAP2’s polymerase activity dominated and no antagonistic effect of TPX2 on growth rate was observed compared to CKAP2 alone (Fig. 5B-D). Together, these findings demonstrate that CKAP2 phosphorylation fine-tunes microtubule polymerization to control spindle size and stability, and that CKAP2 and TPX2 act through overlapping but non-synergistic mechanisms to regulate spindle microtubule dynamics^32,36,46–48^.

## Discussion

Through unbiased proteomic analysis, we identified numerous proteins that associate with CKAP2 during mitosis (Fig. 1; Supplemental Table 1), including prominent MAPs such as Kif2A and TPX2. The presence of multiple chromatin-associated factors, such as SMC2, NCAPG, and TOP2A suggests that these proteins could serve as receptors for CKAP2 during its relocalization to chromatin in late mitosis. Among kinases, AurKA and its cofactor TPX2 were the only candidates detected, consistent with a model in which AurKA-TPX2 represents a potential regulatory axis controlling CKAP2 function during mitosis.

We propose a model in which the AurKA-TPX2 complex phosphorylates CKAP2, thereby constraining spindle growth via a negative feedback loop in which AurKA activity at the spindle initially promotes microtubule growth but, as the spindle matures, CKAP2 phosphorylation feeds back to dampen polymerization, stabilizing spindle size and architecture. CKAP2 is a potent promoter of microtubule nucleation and stabilization; thus, unrestrained activity would risk excessive spindle elongation or misregulation of microtubule mass. Phosphorylation introduces a negative charge, which can weaken electrostatic interactions between CKAP2, which has a pKI of 9.3, and the negatively charged microtubule lattice. Consistent with this, AurKA-dependent phosphorylation of CKAP2 reduces its microtubule-binding affinity *in vitro* and in cells (Figs. 3 and 6G).

Although the structure of CKAP2 on microtubules remains unknown, its regulation may parallel aspects of TPX2 function. TPX2 binds microtubules specifically during mitosis via two helical regions spanning longitudinal and lateral interfaces ^28,32^, whereas CKAP2 associates with microtubules starting in G2^8^. Notably, CKAP2 contains a highly disordered N-terminal region capable of adopting multiple conformations, which could enable flexible binding modes to microtubules and facilitate AurKA access to phosphorylation sites^45^. Indeed, our pull-down experiments revealed that TPX2 interacts most strongly with the N-terminal helix and disordered region of CKAP2 (Fig. 4E), and the presence of TPX2 greatly enhanced AurKA-dependent phosphorylation of CKAP2 *in vitro* (Fig. 4D). We therefore propose that TPX2 acts as both a scaffold and an activator, positioning CKAP2 for efficient phosphorylation by AurKA. Moreover, the addition of microtubules to a kinase reaction containing AurKA and TPX2 potently enhances CKAP2 phosphorylation. We suggest that microtubules promote this reaction by reducing the dimensionality of substrate search, constraining AurKA, TPX2, and CKAP2 from three-dimensional diffusion in solution to a shared two-dimensional lattice. Such lattice-based co-concentration increases effective local abundance and accelerates kinase–substrate encounters, analogous to the mechanism described for AurKB on microtubules^50^.

We identified up to five AurKA consensus sites in the N-terminus of CKAP2 (Fig. 2C, 5A). Among these, Thr39 (mouse Thr52) exhibits the most canonical consensus sequence, though its helical context may limit accessibility. This site has previously been reported in an Aurora B substrate screen^51^, but given the distinct localization of AurKA and Aurora B during mitosis (Fig. S1B-C), direct phosphorylation of CKAP2 by Aurora B in cells is unlikely. Instead, our biochemical, localization, and functional data support AurKA as the primary kinase regulating CKAP2. Thr76 (mouse Thr86) and Thr209 (mouse Thr206) also conform to AurKA consensus motifs. Future structural and biophysical studies will be necessary to elucidate how phosphorylation at this position alters CKAP2’s interaction with microtubules.

In summary, our study identifies CKAP2 as a direct substrate of the AurKA-TPX2 complex and establishes a mechanistic link between spindle-associated kinase signalling and the regulation of microtubule polymerase activity. The CKAP2 interactome presented here provides a valuable foundation for future work to define additional CKAP2 functions and binding partners, both during mitosis and in interphase, and to explore how dysregulation of this pathway may contribute to chromosomal instability in cancer.

## Resource availability

The cell biology datasets generated during and/or analyzed during the current study are available from the corresponding author on reasonable request. The original datasets for CKAP2 interactome have been deposited to the ProteomeXchange Consortium via the PRIDE^52^ partner repository, with the dataset identifiers to be determined following acceptance of the article for publication.

## Acknowledgements

We thank Arnold Hayer and Jose Teodoro for providing reagents. We thank Denis Faubert and Josée Champagne for assistance with the proteomics analysis. We also thank members of the Bechstedt laboratory and Gary Brouhard for helpful suggestions and feedback. The laboratory of SB is supported by grants from the Canadian Institutes of Health Research (CIHR PJT-189995), the Natural Sciences and Engineering Research Council of Canada (NSERC; RGPIN-2024-05603), and in part by a Fonds de Recherche du Québec (Health Sector) Research Centres Grant #288558.

## Author Contributions

Conceptualization: SB and TJK. Investigation: TJK, TL. Formal analysis: TJK, TL. Methodology: TJK, SB. Funding acquisition: SB. Supervision: SB. Writing- original draft: TJK, SB. Writing- review & editing: TJK, SB. Project administration: SB.

## Materials and Methods

### Cell Culture

hTert RPE1 were a gift from Arnold Hayer (McGill University). RPE1 CKAP2 GFP-CKAP2 and CKAP2 knockout cells were described previously^8^. All cells were grown at 37°C in a humidified environment with 5% CO_2_. HeLa cells and U20S cells were purchased from the ATCC and grown in Dulbecco’s modified Eagle medium (Life Technologies #11995073) containing 10% Fetal bovine serum (Wisent #098150), 50 U/mL penicillin and 50 μg/mL streptomycin (Life Technologies #15140122). RPE1 cells were grown in DMEM-F12 (Life Technologies #11320082) with the same supplements. Cells were verified to be free of mycoplasma by frequent staining of plated cells with DAPI. No contamination was observed.

### Inhibitors and reagents

STLC (Fisher Scientific #T25225G) was used at 25 μM. Okadaic acid (Cell Signaling Technology #5934) was used at 100 nM. MLN8237 (Selleck #1028486-01-2) was used at 25-500 nM. ZM447439 (Sigma-Aldrich #189410-5MG). Nocodazole (MilliporeSigma #487929-10MG-M) was used at 100 ng/ml. Palbociclib (MilliporeSigma #PZ0383-5MG) was used at 1 μM. Paclitaxel (Fisher Scientific #AAJ62734MC) was used at varying concentrations. Puromycin (BioBasic # PJ593) was used at 1 μg/ml.

### Plasmids and cloning

Mouse CKAP2 and codon-optimized mouse TPX2 were cloned into pHAT His-USER-GFP-Strep via USER cloning^53^. mmCKAP2 fragments were described previously^45^. The DNA constructs were then verified by Sanger sequencing.

### Transfections

RPE1 or HeLa cells were transfected using Xfect^TM^ transfection reagent (Takara Bio #631318) according to the manufacturer’s directions. Briefly, media on the cells was changed for 500 μl for a 12-well plate. 4 μg of DNA was mixed with 100 μl of buffer. 1.2 μl of Xfect reagent was then added, and the solution mixed by vortex. After 5 minutes incubation, 50 μl of transfection mix was added to cells overnight. The media was then changed 24 hours later.

### Immunofluorescence

Following treatments, cells were fixed in 4% paraformaldehyde in PBS at room temperature for 20 minutes. The cells were washed twice with PBS and then permeabilized in PHEM buffer (60 mM PIPES, 25 mM HEPES, pH 6.9, 10 mM EGTA, and 4 mM MgSO_4_, 1% Triton X-100, 10 mM glycerol 2-phosphate) for 10 minutes. The cells were then washed twice with PBS before blocking with ADB (10% serum, 0.1% Triton X-100 in PBS) for 15 minutes. Primary antibodies were then diluted in ADB and used at the indicated dilution. Secondary antibodies were diluted in ADB. After incubation with antibodies, the cells were washed three times with PBS 0.1% Triton X-100. Finally, the cells were counterstained with DAPI and mounted on slides using FluorSave^TM^ antifade reagent (Millipore #345789-20 ml).

### Immunoprecipitation

For protein-protein interactions, 500,000 RPE1 cells were plated on 10 cm dishes (5 per condition). 24 hours later, palbociclib was added. 18 hours following palbociclib addition, it was washed out with two 4 ml washes with PBS. STLC was then added 5 hours later. 20 hours post-release, the cells were harvested by scraping, washed once with cold PBS, and lysed in the following buffer: 0.5% NP40, 100 mM NaCl, 50 mM Tris pH 7.4, 10 mM glycerol 2-phosphate. Cells were lysed on ice for 20 minutes before centrifuging at 15,000 x g for 20 minutes at 4°C. The supernatant was transferred to a fresh tube, and input sample set aside. 10 μl of packed volume protein G beads were added to each sample together with 5 μg of antibody for 2 hours on a rotating platform at 4°C. The beads were then washed 5 times in lysis buffer and dried using a 30 g needle attached to a vacuum line. 30-100 μl of 1X sample buffer were added to the beads before elution by boiling and loading on gel.

For phosphorylation analysis, cells were prepared and synchronized as above. Following harvesting, the cells were lysed in 50 mM Tris pH 7.5 and 1% SDS. The lysates were then boiled for 5 minutes. SDS was then inactivated by 10-fold dilution with 1.5% Triton X-100 in TBS. Debris was then pelleted via centrifugation at 15,000 x g for 20 minutes at 4°C. CKAP2 was then immunoprecipitated using anti-CKAP2 and protein G agarose as above.

### Mass Spectrometry

CKAP2 was purified from palbociclib-STLC synchronized cells by immunoprecipitation as described above. The entire reaction was then eluted in sample buffer and separated on a 10% SDS-PAGE gel. The gel was then stained with Coomassie blue, and each lane was cut into 12 slices. Gel bands were excised under a clean bench and cut into 1 mm³ pieces. For all subsequent steps, reagent volumes were adjusted according to the volume of the gel pieces. Gel pieces were first washed with water for 5 minutes, then destained twice using a destaining buffer composed of 50 mM ammonium bicarbonate and 50 µL of acetonitrile, each for 10 minutes. Following this, the gel pieces were dehydrated with acetonitrile. Proteins were reduced by incubating the gel pieces in reduction buffer (10 mM DTT, 100 mM ammonium bicarbonate) for 30 minutes at 40 °C and subsequently alkylated with alkylation buffer (55 mM iodoacetamide, 100 mM ammonium bicarbonate) for 20 minutes at 40 °C in the dark. The gel pieces were then dehydrated and washed at 40 °C with acetonitrile for 5 minutes before removing all reagents. They were dried for 5 minutes at 40 °C and then rehydrated at 4 °C for 40 minutes with trypsin solution (6 ng/µL sequencing-grade trypsin from Promega in 25 mM ammonium bicarbonate). The low concentration of trypsin was used to minimize signal suppression and background from autolysis products during LC-MS/MS analysis. Protein digestion was carried out at 58 °C for 1 hour and quenched with 15 µL of 1% formic acid / 2% acetonitrile. The supernatant was transferred to a 96-well plate. Peptide extraction was performed in two 30-minute steps at room temperature using an extraction buffer (1% formic acid / 50% acetonitrile). All peptide extracts were completely dried using a vacuum centrifuge. The LC column used was a PicoFrit fused silica capillary column (15 cm × 75 µm i.d.; New Objective, Woburn, MA), self-packed with C18 reversed-phase material (Jupiter, 5 µm particles, 300 Å pore size; Phenomenex, Torrance, CA) using a high-pressure packing cell. This column was installed on the Easy-nLC II system (Proxeon Biosystems, Odense, Denmark) and coupled to a Q Exactive mass spectrometer (ThermoFisher Scientific, Bremen, Germany) equipped with a Proxeon nanoelectrospray Flex ion source. The chromatography buffers were 0.2% formic acid in water (buffer A) and 100% acetonitrile with 0.2% formic acid (buffer B). Peptides were loaded on-column and eluted using a three-step gradient at a flow rate of 250 nL/min. Solvent B increased from 2% to 35% over 15 minutes, then from 35% to 50% over 7 minutes, and finally from 50% to 90% over 5 minutes. LC-MS/MS data were acquired using a data-dependent Top12 method combined with a dynamic exclusion window of 7 seconds. The mass resolution for full MS scans was set to 70,000 (at m/z 400), and lock masses were used to improve mass accuracy. The mass range for MS scanning was from 360 to 2000 m/z, with a target value of 1 × 10⁶ and a maximum ion fill time of 100 ms. Data-dependent MS2 scan events were acquired at a resolution of 17,500, with a maximum ion fill time of 50 ms, a target value of 1 × 10⁵, an intensity threshold of 1.0 × 10⁴, and an underfill ratio of 0.5%. The normalized collision energy was set to 27%, and the capillary temperature was maintained at 250 °C. Nanospray and S-lens voltages were set to 1.3-1.7 kV and 50 V, respectively. Protein database searches were performed using Mascot 2.6 (Matrix Science) against the UniProt human database. The mass tolerance was set to 10 ppm for precursor ions and 0.5 Da for fragment ions. Trypsin was specified as the enzyme, allowing up to two missed cleavages. Carbamidomethylation of cysteine was set as a fixed modification, and methionine oxidation was included as a variable modification.

### SDS-PAGE and Western blot

Gels and blots were performed as described previously ^54^. Prior to loading samples on gel, 4X Laemmli buffer (200 mM Tris pH 6.8, 4% SDS, 40% glycerol, 4% 2-mercaptoethanol, 0.12 mg/ml bromophenol blue) was added to a final dilution of 1X, and the samples were boiled for 5 minutes. Proteins were separated on SDS-PAGE gel using stacking gel (4% 29:1 acrylamide: Bis-acrylamide, 125mM Tris pH 6.8, 0.1% SDS, 0.1% ammonium persulfate, 0.1% TEMED) and (8-15% 29:1 acrylamide: Bis-acrylamide, 400 mM Tris pH 8.8, 0.1% SDS, 0.1% ammonium persulfate, 0.1% TEMED) at 120V until the bromophenol blue has run off or longer, as needed. Transfer onto nitrocellulose membrane was performed for at least 24 hours at 30V under wet conditions in 1X transfer buffer (14.4 g/L glycine, 3.0 g/L Tris, 20% methanol). Conditions for western blots include the use of 5% nonfat dry milk in TBS-T 0.5% (50 mM Tris [pH 7.2], 150 mM NaCl, 0.5% Tween 20) for blocking and TBS-T 0.5% for washing. 4 washes for 10 minutes each were performed after primary and secondary antibody incubation periods. The bands were visualized by enhanced chemiluminescence using Clarity (Bio-Rad #1705060) using a Bio-Rad ChemiDoc^TM^ Touch Imaging System.

### *In vitro* analysis of CKAP2

Mouse CKAP2 protein variants were produced in the *E. Coli* strain BL21(DE3). The bacteria were suspended in 25 ml buffer A (50 mM NaPO_4_, 10 mM Imidazole, 300 mM NaCl, 10% Glycerol, pH 8) and processed in an Emulsiflex high-pressure homogenizer (Avestin). The lysate was then ultracentrifuged at 45Ti rotor at 25000 RPM for 45 minutes. 200 μl of packed glutathione agarose was then added for 2 hours on a rotating platform. 1 ml Ni-NTA agarose was then added to the lysate for 2 hours on a rotating platform. The agarose was then washed, and purified proteins eluted with buffer A containing 300mM imidazole and then dialyzed against 0.4X TBS. Glycerol was then added to 10% and the proteins frozen until further use. Kinase reactions were performed by mixing CKAP2, GFP-AurKA-6XHis, 100 μM ATP in kinase buffer 50 mM Tris pH 7.2, 25 mM NaCl, 10mM MgCl_2_ 1mM DTT, 0.01% Triton X-100 for 1 hour at 37°C. The reactions were stopped by addition of 4X SDS sample buffer, and half the reaction loaded on SDS-PAGE gel. The gels were then stained with ProQ Diamond phosphoprotein stain (Thermo Scientific) according to the manufacturer’s instructions and subsequently with Coomassie blue.

*In vitro* microtubule (MT) assays were performed as described previously^9^. CKAP2 fragments were produced in BL21(DE3) *E. coli* grown in LB media to OD_600_ and then induced with 0.5 mM Isopropyl-D-1-thiogalactopyranoside (IPTG) at 20°C overnight. The bacteria were then harvested and lysed in Buffer A (50 mM NaH2PO4, 300 mM NaCl, 0 mM imidazole, pH 7.8) using an EmulsiFlex-C5 (Avestin Inc, Canada). Debris was then pelleted by ultracentrifugation at 30,000 x g for 1 hour at 4°C. The lysate was then loaded onto a column containing His60 Ni SuperFlow Resin (Takara Bio, USA). Purified protein was eluted with Buffer B (50 mM NaH2PO4, 300 mM NaCl, 200 mM imidazole, pH 7.8). Eluted proteins were then loaded onto a column containing Strep-Tactin Sepharose resin (IBA Lifesciences, Germany), washed with 5 column volumes of Buffer W (100 mM Tris-HCl, 150 mM NaCl, 1 mM EDTA, pH 7.8) and finally eluted with Buffer W supplemented with 2.5 mM Desthiobiotin. Protein concentration was determined by measuring the absorbance at 280 or 488 nm using a DS- 11 FX spectrophotometer (DeNovix Inc, USA). SDS-PAGE was used to determine the purity of the proteins. Tubulin was purified from bovine brains as previously described^55^ with the difference of having used Fractogel EMD SO3- (M) resin (Millipore-Sigma) instead of phosphocellulose. Tubulin was labelled using CF640R-NHS-Ester (Biotium, USA) and tetramethylrhodamine (TAMRA, Invitrogen Inc, Canada), as previously described^56^. Before use, an additional cycle of MT polymerization was performed.

### Microscope Slide Channel Preparation

Glass coverslips (22×22 and 18×18 mm) were washed with acetone and then sonicated in 50% methanol with 0.5 M KOH and finally rinsed with water. The coverslips were then dried and exposed to air plasma (Plasma Etch) for 3 minutes before silanizing by soaking them in 0.1% dichlorodimethylsilane in *n*-heptane, before sonicating them in *n*-heptane and finally in ethanol. Flow channels with a volume of 7 μl were created by using two pieces of silanized coverslips held by double-sided tape in custom coverslip holders. The channels were incubated with anti-TAMRA antibodies for 5 minutes, blocked with 5% Pluronic F-127 for 10 minutes and washed with BRB80 before incubation with MTs. The channels were used at 35 ^°^C using a heated microscope objective and an incubation chamber.

### Binding to Paclitaxel-stabilized MTs

GDP-MTs stabilized by paclitaxel were prepared by polymerizing a 1:15 ratio of TAMRA-labelled to unlabeled tubulin in the presence of 4 mM MgCl_2_, 1mM GTP, 5% DMSO in BRB80 buffer. The mixture was then incubated for 30 minutes at 37 ^°^C. The MTs were then diluted using BRB80 plus 10 μM paclitaxel, pelleted at 100,000 RPM at 30 psi for 7 minutes using an Airfuge (Beckman-Coulter) and finally resuspended in BRB80 plus μM paclitaxel.

To assess binding of CKAP2 to MTs, A combination of GMPCPP double-cycled MTs and paclitaxel-stabilized MTs was mixed and flowed into microscope slide channels. Since the paclitaxel-stabilized MTs contain less TAMRA label, they possess a dimmer signal compared to GMPCPP MTs.

### Dynamic Microtubule Assay

GMPCPP MT seeds were created by polymerizing a 1:4 ratio of TAMRA-labelled to unlabeled tubulin in the presence of 1 mM guanosine-5’- [(α, β)-methyleno] triphosphate (GMPCPP, Jena Biosciences, Germany), a non-hydrolyzable GTP analogue during two cycles. On the day of an experiment, unlabeled and CF642R-labeled tubulin were mixed at a 1:17 ratio, aliquoted and flash-frozen in liquid nitrogen. To start an experiment, GMPCPP MT seeds were flowed into a channel and then washed with BRB80 buffer. MT growth was initiated by introducing tubulin (8 μM) in BRB80 buffer containing 1 mM GTP, 40 mM D-glucose, 64 nM catalase, 250 nM glucose oxidase, 10 mM DTT, 0.1 mg/ml BSA together with CKAP2. Images were then taken at suitable time points to track MT growth and catastrophe.

### Microscopy

High magnification images were acquired with either a Zeiss Axio Observer 7 fitted with a TIRF equipped with a Prime 95B CMOS camera (Photometrics) with a pixel size of 107 nm controlled by Zeiss Zen 2.5 Blue Edition software version or a Nikon Eclipse Ti microscope spinning disc confocal (Cicero; Crest Optics) microscope equipped with AURA III Light Engine (Lumencor®) and Hamamatsu ORCA-fusion C14440 camera with 110 nm pixel size controlled by Nikon-Elements AR 6.10.01 software. Images were acquired in 0.15-0.4 μm sections using a plan apo 1.4 numerical aperture 100X or 60X oil-immersion objective using 1X1 binning. All image analysis, adjustments, measurements and cropping were performed using Fiji software ^57^. Widefield Image deconvolution was performed using Zen Blue Edition 2.5 software. All protein intensity analysis was performed on the raw images. Line scans were performed on the deconvoluted images in Fiji and the data was then exported to Graphpad Prism version 8.4.3. or version 10.2.1.

### Antibodies

The following antibodies were used for immunofluorescence (IF) and/or immunoblotting (IB): mouse anti-AurKA (Cell Signaling Technology 1F8; IF at1:1000, blot at 1:500), Rabbit anti-AurKA pT288 (Cell Signaling Technology C39D8; IF at 1:500), mouse anti-TPX2 (Cell Signaling Technology D9Y1V; IF at 1:1000, blot at 1:500), rabbit anti-Aurora B (Cell Signaling Technology 3094T; IF at 1:50), mouse anti- β−Tubulin (Sigma clone 2.1 T4026; IF at 1:200), rabbit anti-CKAP2 (Proteintech 25486-1-AP; blot at 1:500), Rabbit anti-CKAP2 (AbClonal A9706; IF at 1:200), Mouse anti-TAMRA (Invitrogen 5G5; label at 1:100). Secondary antibodies used were highly cross-adsorbed Alexa Fluor® 488, 594 and 647 raised in donkey or goat (Invitrogen; IF at 1:1000), horseradish peroxidase (Bio-Rad; IB at 1:10000-10,0000), Anti-IgG (H+L) Goat Polyclonal Antibody (Horseradish Peroxidase), Peroxidase AffiniPure Goat Anti-Rabbit IgG (H+L) (IB at 1:5000), Goat Polyclonal Antibody (Horseradish Peroxidase), Peroxidase AffiniPure Goat Anti-Mouse IgG (H+L) (IB at 1:5000).

### Structural Models

Structural models of the interaction of CKAP2 (Uniprot accession #Q3V1H1) with TPX2 (#A2APB8) have been generated using AlphaFold2.

### Statistical analysis of cell biology and microtubule in vitro data

All statistical tests were performed using Graphpad Prism version 8.4.2. Except where indicated, all experiments were performed as 2-3 independent biological repeats. In most IF experiments, 25 cells for each condition were measured. Experiments with other numbers are noted in the figure legends.

---

**Figure S1:**
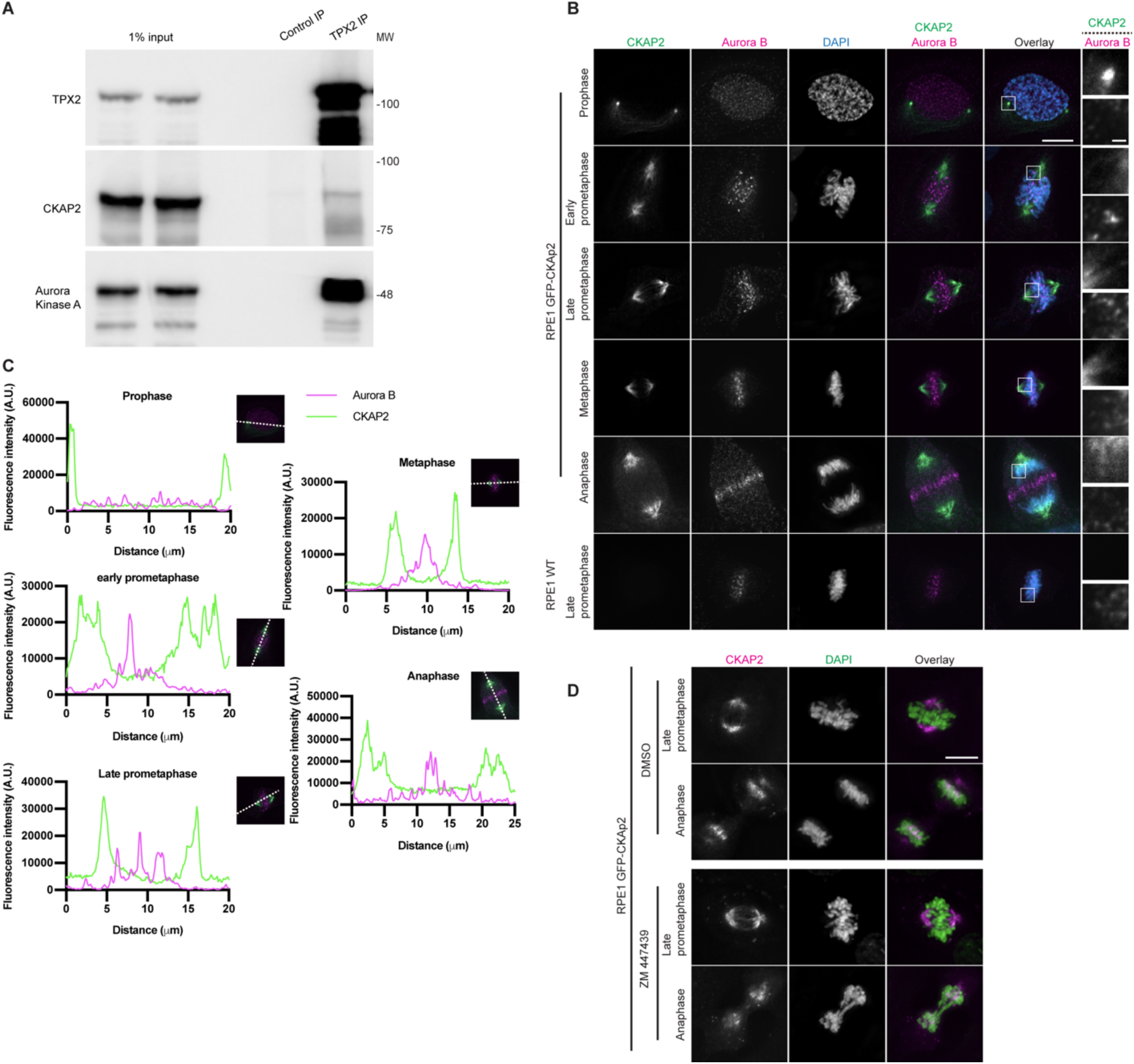
AurKB does not co-localize with CKAP2 during mitosis; TPX2 is a CKAP2 binding partner. A) TPX2 was immunoprecipitated together with interacting proteins from RPE1 cells synchronized via Palbociclib-STLC block. The purified proteins were then separated by SDS-PAGE, transferred to a nitrocellulose membrane and blotted as indicated. The panels were adjusted for brightness and contrast. An experiment representative of two independent biological repeats is shown. B) Immunofluorescence images from RPE1 eGFP-CKAP2 cells in various stages of mitosis stained for CKAP2, AurKB, and DAPI. The CKAP2 and AurKB channels were adjusted evenly for brightness and contrast. The DAPI channel of each condition was adjusted independently. Representative images from 2 independent biological experiments are shown. Scale bars, 10 μm and 1 μm (inset). C) Line scans across the indicated paths from the images in panel (B). D) RPE1 GFP-CKAP2 cells were treated with 2 μM ZM447439 for 1 hour and then fixed and stained with DAPI. The CKAP2 channel was adjusted evenly for brightness and contrast. The DAPI channel was adjusted independently. Representative images from 2 independent biological experiments are shown. Scale bar, 10 μm. E) 1L cultures of BL21 E. Coli expressing the indicated contstructs were induced with 0.75 mM IPTG overnight at 18°C. The bacteria were then harvested by centrifugation and lysed using an Emulsiflex homogenizer. The bacterial lysates were then ultracentrifuged to separate cell debris. The lysates were then added to columns containing His60 Ni SuperFlow Resin (CKAP2 and TPX2 only). The columns were washed and purified proteins were then eluted and purified again using columns containing Strep-Tactin Sepharose resin. The columns were washed and then the purified proteins eluted. 1 μg of purified protein was migrated on SDS-PAGE gel, which was then stained with Coomassie. A representative image of at least 2 independent experiments is shown. The image was adjusted for brightness and contrast for presentation.

**Supplementary Movie 1:** AlphaFold2 model with CKAP2 domains colour-coded as in Figure 4 and TPX2 in gold. Zoom into the N-terminal TPX2 alpha helix interacting with CKAP2.

